# A genome-wide CRISPR screen in human prostate cancer cells reveals drivers of macrophage-mediated cell killing and positions AR as a tumor-intrinsic immunomodulator

**DOI:** 10.1101/2023.06.06.543873

**Authors:** Anniek Zaalberg, Emma Minnee, Isabel Mayayo-Peralta, Karianne Schuurman, Sebastian Gregoricchio, Thijs A. van Schaik, Liesbeth Hoekman, Dapei Li, Eva Corey, Hans Janssen, Cor Lieftink, Stefan Prekovic, Maarten Altelaar, Peter S. Nelson, Roderick L. Beijersbergen, Wilbert Zwart, Andries Bergman

## Abstract

The crosstalk between prostate cancer (PCa) cells and the tumor microenvironment plays a pivotal role in disease progression and metastasis and could provide novel opportunities for patient treatment. Macrophages are the most abundant immune cells in the prostate tumor microenvironment (TME) and are capable of killing tumor cells. To identify genes in the tumor cells that are critical for macrophage-mediated killing, we performed a genome-wide co-culture CRISPR screen and identified AR, PRKCD, and multiple components of the NF-κB pathway as hits, whose expression in the tumor cell are essential for being targeted and killed by macrophages. These data position AR signaling as an immunomodulator, and confirmed by androgen-deprivation experiments, that rendered hormone-deprived tumor cells resistant to macrophage-mediated killing. Proteomic analyses showed a downregulation of oxidative phosphorylation in the *PRKCD-* and *IKBKG-KO* cells compared to the control, suggesting impaired mitochondrial function, which was confirmed by electron microscopy analyses. Furthermore, phosphoproteomic analyses revealed that all hits impaired ferroptosis signaling, which was validated transcriptionally using samples from a neoadjuvant clinical trial with the AR-inhibitor enzalutamide.

Collectively, our data demonstrate that AR functions together with the PRKCD and the NF-κB pathway to evade macrophage-mediated killing. As hormonal intervention represents the mainstay therapy for treatment of prostate cancer patients, our findings may have direct implications and provide a plausible explanation for the clinically observed persistence of tumor cells despite androgen deprivation therapy.

## Introduction

Prostate cancer (PCa) is the second most common malignancy in men^1^. While surgery and radiotherapy remain the main treatment options for localized PCa, patients presenting with advanced or metastatic stages of disease cannot be cured. Therefore, alternative therapeutic options are of utmost clinical relevance to improve treatment and survival of these patients. The prostate tumor microenvironment (TME) plays a critical role in the development and progression of PCa and altering the TME could potentially provide novel ways to treat patients and improve survival^2, 3^. In particular, the inflammatory microenvironment consisting of predominantly tumor-associated macrophages (TAMs) is involved in multiple opposing processes and represents an essential modulator of malignant progression, metastases and the overall therapeutic response^4^. The rate of TAM infiltration in PCa is associated with disease progression after hormonal therapy and preclinical studies have suggested that TAMs modulate PCa cell proliferation and migration^5, 6^. A large spectrum of TAM phenotypes have been described, ranging from classically activated, pro-inflammatory and anti-tumor M1 macrophages to anti-inflammatory and pro-tumor M2 macrophages^7^. Theoretically, M1 macrophages can prevent cancer development and progression, but apparently their anti-tumor function is impaired since cancer develops in their presence^8, 9^. Therefore, means to enhance the cell-killing by M1 macrophages could provide potential therapeutic benefits.

Androgen receptor (AR) signaling plays a key role in normal prostate physiology, as well as in the development and progression of PCa^10^. AR is a hormone-driven transcription factor that is activated upon androgen stimulation whereafter it drives transcription of androgen responsive genes, essential for cell proliferation and survival^11^. Among its involvement in multiple cellular processes, AR has been previously identified by us and others as a modulator of innate and adaptive immune cells, including macrophages and T-cells^12–14^. AR inhibition is the single most effective treatment for metastatic PCa. However, although most patients initially benefit from AR signaling inhibition therapy, treatment resistance inevitably occurs, leading to the development of lethal castration resistant PCa (CRPC).

To date, the interplay between PCa cells and macrophages is poorly understood and the critical tumor-intrinsic features required for macrophage-mediated tumor killing remain unknown. In this study, we performed a comprehensive genome-wide CRISPR screen to identify genes in the tumor cell, that are required for macrophage-mediated killing. These studies identified protein kinase C delta (PRKCD) and several components of the NF-κB pathway as critical mediators. Strikingly, AR was identified as a critical hit, essential for tumor cell killing by macrophages, and hormone-deprivation of the tumor cells prevented them from being killed by M1 macrophages. These data position AR as a *bona fide* tumor cell-intrinsic immune modulator and reveal immune protection from macrophages as an adverse consequence of hormonal therapy in PCa patients.

## Results

### A genome-wide co-culture CRISPR screen identifies essential drivers of macrophage-mediated prostate cancer cell killing

To study the interaction between PCa cells and pro-inflammatory M1 macrophages, AR responsive LNCaP cells were co-cultured with THP-1 derived macrophages. Tumor cell proliferation capacity was greatly reduced in the presence of M1 macrophages (co-culture) as compared to PCa cells alone (monoculture) (Figure 1A). In agreement with this observation, tumor cell viability was also markedly decreased in co-culture conditions compared to monoculture control (Figure 1B). Based on these observations, we set out to identify which genes in PCa cells are essential for macrophage-mediated killing, by performing a genome-wide co-culture CRISPR screen in LNCaP cells. Tumor cells were transduced with the Brunello library, consisting of 77,441 single guide RNAs (sgRNAs), with an average of 4 sgRNAs per gene and 1000 non-targeting control sgRNAs^15, 16^. Tumor cells were cultured with (co-culture) or without (T0) macrophages, to identify genes that, when knocked out, were individually capable of conferring resistance to macrophage-mediated killing (Figure 1C). As a quality control, practically the entire Brunello library was identified in the starting population of the screen (T0) (77,041 sgRNAs in the library, 76,994 sgRNAs in our data) (Supplementary Figure 1A). To compare counts between the samples, a relative factor based on the total counts of each sample was calculated and used to normalize the count values (Supplementary Figure 1B). Based on a significance cut-off of an FDR < 0.1 and a Log2FCMedian of > 1.5 we identified sgRNAs for 6 genes, enriched in the tumor/macrophage co-culture arm compared to T0. These genes when knocked out prevented tumor cell death in the co-culture population, and include: *IKBKB, IKBKG, CHUK, PRKCD* and *AR* (Figure 1D). These data suggest a critical role for these genes in regulating macrophage-mediated killing in PCa cells. IKBKB, IKBKG and CHUK are part of the IKK kinase complex, the central regulator of NF-κB signaling^17^. Protein kinase C delta (PKCδ) is a serine/threonine kinase that has been shown to both positively and negatively regulate cell-cycle progression^18, 19^ and plays a dual role in cell death^18^. It has been shown that upon DNA damage (e.g., genotoxic stress), PKCδ mediates p53 (in)dependent apoptosis^20^. Surprisingly, AR was a top hit in our CRISPR screen, suggesting that the absence of AR signaling protects PCa cells from M1 macrophage-mediated cytotoxicity, indicating the immunomodulatory capacity of this essential PCa driver.

**Figure 1.**
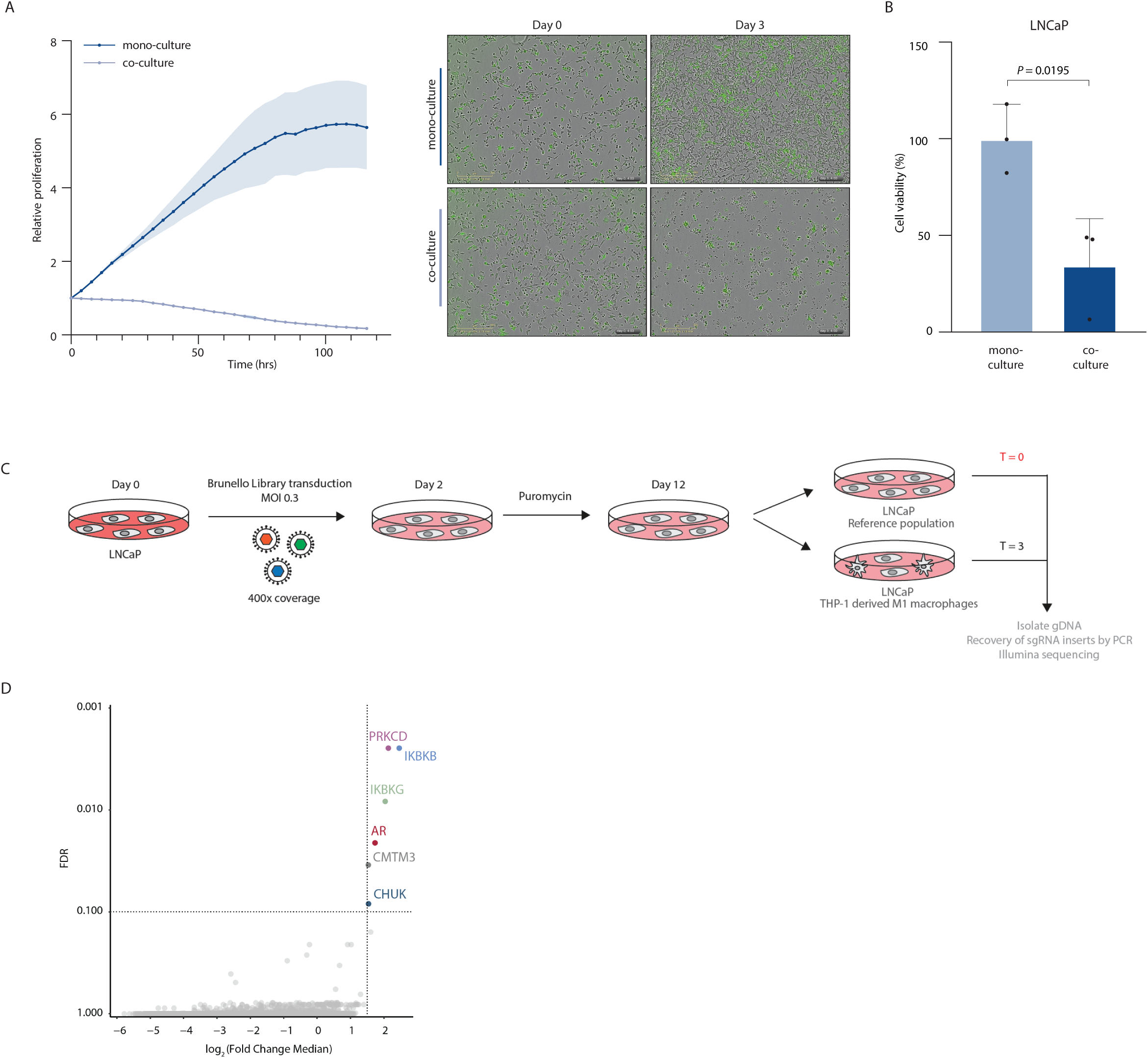
PCa cells are sensitive to macrophage-mediated killing which is driven by NF-κB-, PRKCD-and AR-signaling. A. Cell proliferation of LNCaP-eGFP cells cultured alone (monoculture, dark blue line) or in a co-culture (light blue line) with pro-inflammatory macrophages (n = 6, mean values are depicted and error bars represent the standard deviation). Y-axis depict relative proliferation based on eGFP mean intensity (left side). IncuCyte microscopy images of LNCaP-eGFP monoculture and LNCaP-eGFP-macrophage co-cultures at timepoint 0 (day 0) and after three days of culturing (day 3). Scale bar = 300μm. B. Cell viability assay of LNCaP cells cultured alone (monoculture) or in a co-culture with pro-inflammatory macrophages. Data is normalized to the monoculture condition. Mean values and the standard deviations are depicted (n=3, mean values are depicted and error bars represent the standard deviation). P-value was calculated using a two-tailed unpaired t-test. C. Schematic representation of the genome-wide co-culture CRISPR-screen for the identification of drivers of macrophage-mediated killing. D. Results of the screen. Scatter plot of the log_2_FcMedian (fold change of co-culture arm vs T0) and the false discovery rate (FDR). Dotted lines indicate the threshold (log_2_FcMedian > 1.5 and FDR < 0.1). Analyses were performed using DESeq2 and MaGECK. Colored dots indicate the significant hits of the screen.

### Impaired NF-κB-signaling promotes resistance of macrophage-mediated tumor cell killing

Our co-culture screen identified multiple components of the IKK complex as critical for macrophage-mediated tumor killing. The IKK complex represents the core element of the NF-κB signaling pathway^17^, which consists of two kinases IKKα (CHUK) and IKKβ (IKBKB) and a regulatory subunit IKKγ (IKBKG). In response to stimuli, - such as inflammatory cytokines, infections or other cellular stress (e.g., DNA damage or UV stress) - the NF-κB inhibitor IκB becomes degraded, releasing NF-κB subunits p50 and p65 to activate transcription of NF-κB responsive genes^21^ (Figure 2A). To validate our NF-κB signaling associated CRISPR screen hits, we generated IKBKB, IKBKG and CHUK knock-out (KO) models using two independent sgRNAs per gene. CRISPR-mediated targeting of IKBKB, IKBKG or CHUK resulted in a significantly reduced expression of the target mRNA (Figure 2B) and protein (Figure 2C) level. Knockout of *IKBKB, IKBKG or CHUK* strongly reduced NF-κB activity as measured by luciferase reporter assays (Supplementary Figure 2), confirming the essential function of each individual component in NF-κB signaling.

**Figure 2.**
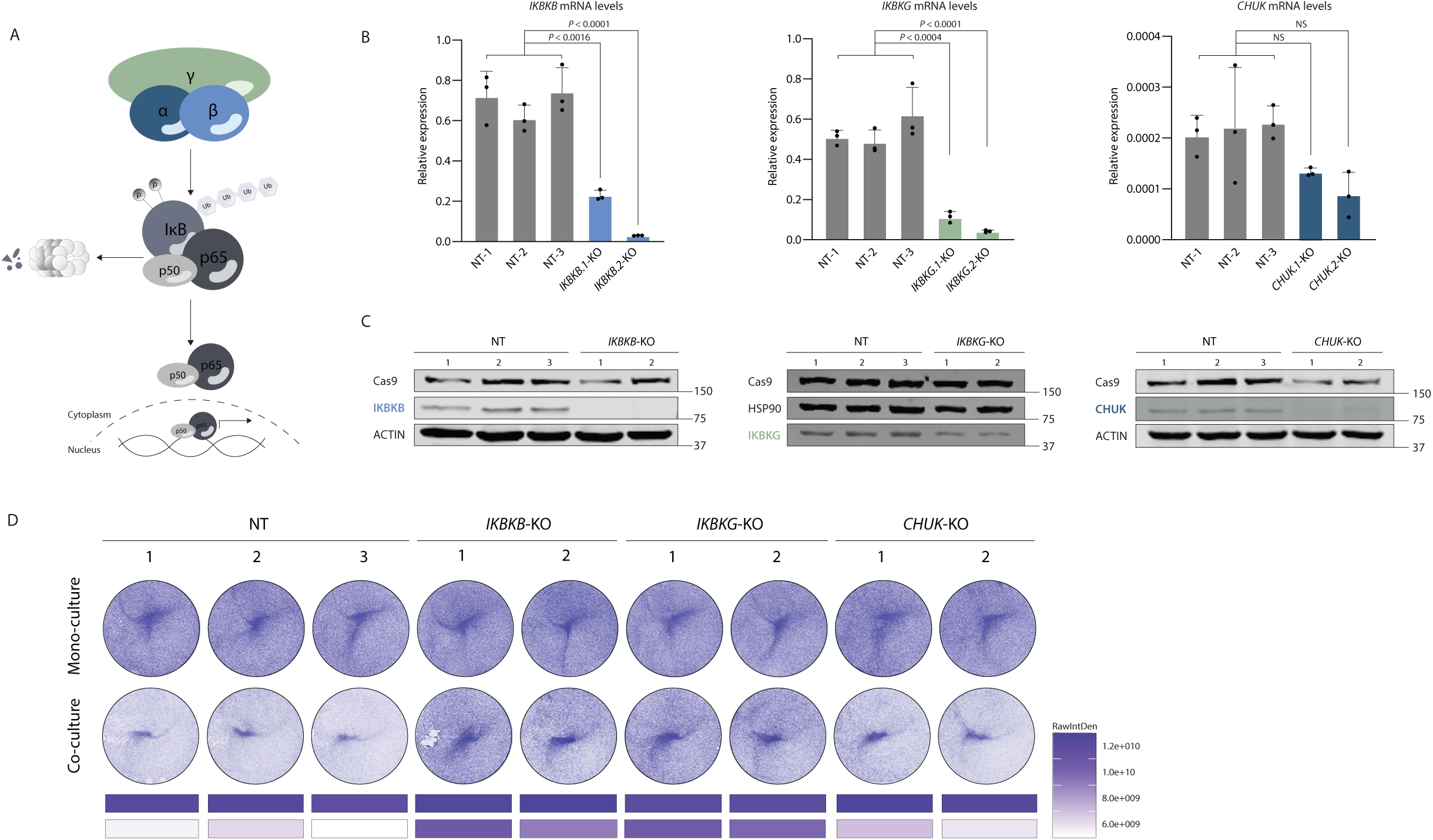
Resistance of PCa cells to macrophage-mediated killing is mediated by members of the IKK complex. A. Schematic overview of IKK signaling. Canonical NF-κB is regulated by the IKK complex, which consists of the IKKα, β, and γ subunits. Phosphorylation of IκBα leads to its degradation, allowing the p65-p50 dimer to accumulate in the nucleus and regulate transcription of target genes. B. Relative mRNA levels of IKBKB, IKBKG and CHUK (normalized to *TBP*) in *IKBKB-KO*, *IKBKG-KO* and *CHUK-KO* LNCaP cells (n=3, mean values are depicted and error bars represent the standard deviation). P-value was calculated using a two-way ANOVA test. C. Protein levels of IKBKB, IKBKG and CHUK protein levels in *IKBKB-KO*, *IKBKG-KO* and CHUK-KO LNCaP cells. Actin and HSP90 are used as loading control. Cas9 was used as a control for efficient lentiCRISPRv2 transfection. D. Crystal violet assay of NT control and *IKBKB-KO*, *IKBKG-KO* and CHUK-KO cells cultured as a monoculture (up) and co-culture (down) with macrophages. Cells were cultured for 3 days. Crystal violet staining was quantified using Fiji software.

Next, all three knockouts were tested to validate our screen results. *CHUK* could not be validated as a hit, as macrophage-mediated cell death was not perturbed, possibly due to incomplete knockout (Figure 2D). However, disruption of either *IKBKB* or *IKBKG* resulted in resistance to macrophage-mediated killing compared to Non-Targeting (NT) control, validating our screen results (Figure 2D). Cumulatively, our data demonstrate that single members of the IKK complex in PCa cells are critical for macrophage-mediated killing, highlighting NF-κB action as a relevant component in the interplay between PCa and its microenvironment.

### Depletion of PKCδ induces tumor resistance to macrophage-mediated killing

The sgRNAs targeting *PRKCD* were among the top hits in our screen, preventing macrophages from effectively killing PCa cells. The PKC family of serine/threonine protein kinases consists of 11 isoforms that regulate a variety of biological pathways^22^. PKCδ is involved in many cellular processes including cell proliferation and survival, apoptosis, signal transduction, transcriptional hormonal regulation and immune responses^23–29^. Interestingly, *PRKCD* is an androgen-responsive gene, as its mRNA and protein levels were increased in LNCaP cells upon synthetic androgen (R1881) treatment in a dose-dependent manner (Figure 3A and 3B), confirming previous studies^30, 31^. To further validate these findings, we focused on the PRKCD locus and evaluated AR chromatin binding (as assessed by ChIP-seq^32^) and chromatin looping to the transcription start site (TSS) of *PRKCD* (as assessed by H3K27ac Hi-ChIP^33, 34^) in LNCaP cells. These analyses revealed several AR binding sites (ARBS) around the PRKCD locus and confirmed enhancer interactions with the TSS of PRKCD (Figure 3C). To validate the dependence of PRKCD in LNCaP cells on macrophage-mediated tumor cell death, we generated PRKCD-KO models using two independent sgRNAs. Loss of *PRKCD* significantly reduced its expression, both on mRNA (Figure 3D) and protein level (Figure 3E). Ultimately, disruption of *PRKCD* resulted in resistance to macrophage-mediated killing compared to NT control (Figure 3F). Taken together, these data indicate that *PRKCD* is an androgen-responsive gene that plays a critical role in facilitating macrophage-mediated cell death.

**Figure 3.**
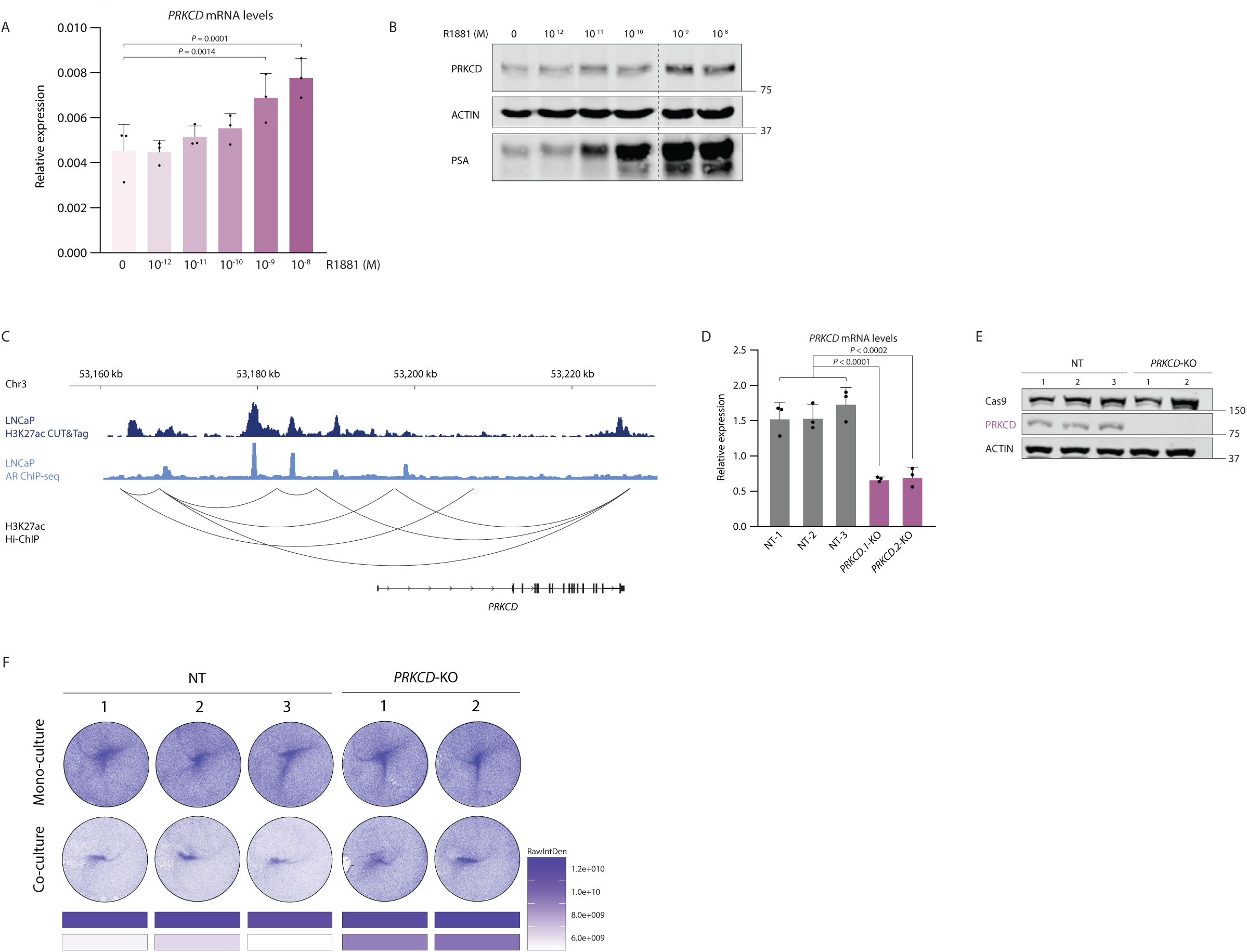
PRKCD is an androgen responsive gene important for regulating macrophage-mediated PCa cell killing. A. Relative mRNA levels of PRKCD (normalized to *ACTIN*) in LNCaP cells stimulated with vehicle (DMSO) or R1881 in a range from 10^-12^-10^-8^M (n=3, mean values are depicted and error bars represent the standard deviation). P-value was calculated using a two-way ANOVA test. B. Protein levels of PKCδ increase in a dose-dependent manner. Protein levels of PKCδ in LNCaP cells stimulated with vehicle (DMSO) or R1881 in a range from 10^-12^-10^-8^M. Actin was used as loading control. C. Genomic snapshot of the enhancer region upstream of the PRKCD locus with H3K27ac Hi-ChIP interaction data. AR ChIP-seq tracks and H3K27ac CUT&Tag are shown. D. Relative mRNA levels of PRKCD (normalized to *TBP*) in *PRKCD*-KO LNCaP cells (n=3, mean values are depicted and error bars represent the standard deviation). P-value was calculated using a two-way ANOVA test. E. Protein levels of PKCδ in *PRKCD*-KO LNCaP cells. Actin is used as loading control. Cas9 was used as a control for efficient lentiCRISPRv2 transfection. E. Crystal violet assay of NT control and PRKCD-KO cells cultured as a monoculture (up) and co-culture (down) with macrophages. Cells were cultured for 3 days. Crystal violet staining was quantified using Fiji software.

### Oxidative phosphorylation is impaired in KO cells and confers resistance to macrophage-mediated killing by ferroptosis

Having identified that the IKK complex and PKCδ are critical for cell death mediated by pro-inflammatory macrophages, we next sought to gain a better understanding of the mechanism underlying the evasion of macrophage-mediated killing capacity. Therefore, we investigated whether there are any proteomic differences in IKBKG-KO and PRKCD-KO cells compared to the control cells (Figure 4A). As expected, both IKBKG and PKCδ proteins were effectively depleted in our KO cell line models as they were strongly depleted in respectively the *IKBKG-KO* and *PRKCD-KO* cells. Interestingly, Gene Set Enrichment Analysis (GSEA) on ranked differentially expressed proteins in both *PRKCD-KO* and *IKBKG-KO* cell lines showed a marked depletion of the oxidative phosphorylation and citric acid cycle pathways (Figure 4B). This was accompanied by a strong significant downregulation of oxidative phosphorylation (Hallmark gene sets; #M5936) at the whole proteome level for both KOs (Figure 4C). Since proteins important for oxidative phosphorylation are embedded in the lipid bilayer of the inner mitochondrial membrane (IMM)^35^, we hypothesized that the mitochondria in our IKBKG-KO and PRKCD-KO cell models are functionally impaired. Therefore, we imaged mitochondrial structures using electron microscopy. In both *IKBKG-KO* and *PRKCD-KO* cells, morphological features of mitochondria were strongly affected, being highly condensed, with irregular cristae and linearization of cristae membranes (Figure 4D). However, given that the ability of cells to proliferate was not affected in either KO (Supplementary Figure 3), we conclude that the core mitochondrial function remained intact. In addition, we performed phospho-proteomic analysis to understand the cellular signaling in our KO cell line models. Interestingly, ingenuity pathway analysis (IPA) on the phospho-proteomes of the *IKBKG-KO* and the *PRKCD-KO* cell lines showed a decrease in ferroptosis signaling compared to NT control cells (Figure 4E). This was observed in FBS conditions as well as upon TNFα stimulation, which is one of the main cytokines secreted by pro-inflammatory macrophages^36^. The role of mitochondria in ferroptosis has been controversial, but recent work has demonstrated that the causal contribution of mitochondrial events such as ROS accumulation and accumulation of lipid peroxidation are the final steps in ferroptosis^37, 38^ . In conclusion, both the IKK complex and PKCδ are critical components in macrophage-mediated PCa cell death, functionally converging on mitochondrial activity and the induction of ferroptosis.

**Figure 4.**
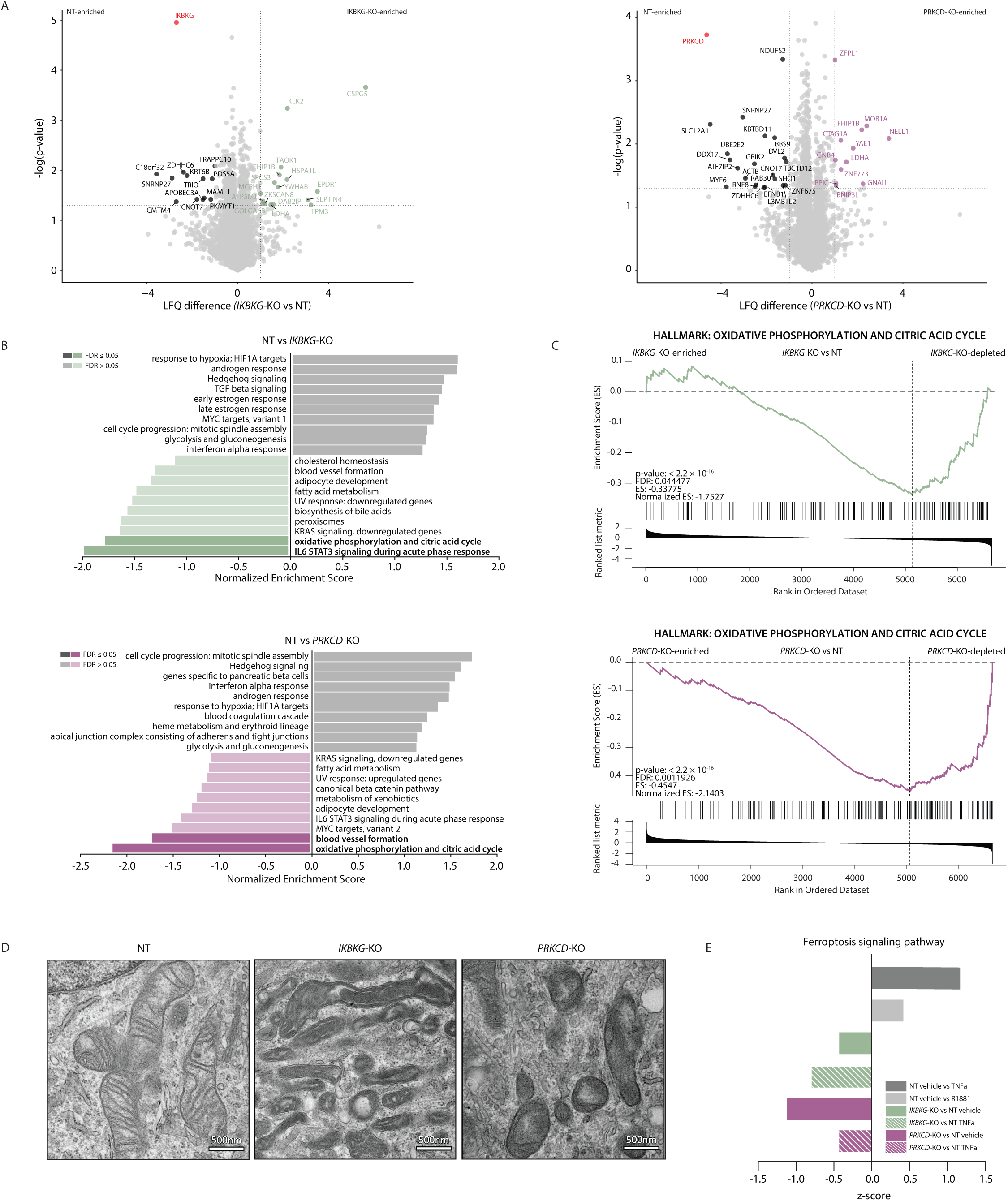
Oxidative phosphorylation is downregulated in *IKBKG-KO* and *PRKCD*-KO leading to phenotypically altered mitochondria and downregulation of ferroptosis signaling. A. Proteomic analysis of non-targeting (NT) control, *IKBKG*-KO and *PRKCD*-KO *LNCaP* cells. Left: volcano plot comparing total proteome of *IKBKG*-KO over NT cells. Proteins highlighted in green are enriched in the *IKBKG*-KO cells, proteins highlighted in black are enriched in the NT control cells. Significance cut-off is depicted with a dotted line (LFQ > 1, -log(p-value) > 1.3) (n=4). Right: volcano plot comparing total proteome of PRKCD-KO over NT cells. Proteins highlighted in purple are enriched in the PRKCD-KO cells, proteins highlighted in black are enriched in the NT control cells. Significance cut-off is depicted with a dotted line (LFQ > 1, -log(p-value) > 1.3) (n=4). B. GSEA enrichment analysis for the hallmark oxidative phosphorylation (M5936) comparing *IKBKG-KO* vs NT (top) and PRKCD-KO vs NT (bottom). Ranking is based on the LFQ values of the proteome data. FDR was determined using WEB-based GEne SeT AnaLysis Toolkit (WebGestalt). C. GSEA enrichment profiles for the hallmark oxidative phosphorylation (M5936) comparing *IKBKG-KO* vs NT (top) and PRKCD-KO vs NT (bottom). Ranking is based on the LFQ values of the proteome data. Nominal p-value and NES were determined with GSEA. D. Electron microscopy images of mitochondrial pellets isolated from NT (left), *IKBKG-KO* (middle) and PRKCD-KO cells (right). Scale bar = 500nm. E. Pathway analysis of the phosphoproteomics data of the *IKBKG-KO*, *PRKCD-KO* and NT control cells treated with vehicle (DMSO), TNFα or R1881 using the Ingenuity Pathway Analysis (IPA) software. The ferroptosis signaling pathway is depicted. Bars display negative or positive z-scores.

### AR acts as an immunomodulator and regulates macrophage-mediated PCa cell killing

Based on our CRISPR screen, we conclude that AR expression in the tumor cells was essential for macrophage-mediated killing. As AR is the critical drug target in PCa care, these results suggest that therapeutic inhibition of AR is involved in tumor cell resistance to killing by macrophages. To test this hypothesis, LNCaP cells were hormone-deprived, and subsequently treated with vehicle control or R1881 to activate AR signaling, after which the co-culture with pro-inflammatory macrophages was initiated. Cell viability was measured after 3 days of co-culture (Figure 5A). Indeed, cells cultured in androgen deprived conditions are resistant to macrophage-mediated killing, whereas R1881 treatment restored tumor cell killing capacity (Figure 5B). These results imply that AR serves as an immunomodulator and that AR activity is critical for susceptibility to macrophage-mediated cell death. To further substantiate these findings, we next performed additional co-culture experiments using LNCaP-abl (androgen ablated) cells with constitutively active AR signaling^34, 35^, and AR-negative PC3 cells (Figure 5C). In line with its AR status, LNCaP-abl cells were susceptible to macrophage-mediated killing, irrespective of the presence of hormone (Figure 5C, middle). In contrast, AR-negative PC3 cells were resistant to macrophage-mediated killing, both in androgen deficient and proficient conditions (Figure 5C, right). Importantly, these results were fully recapitulated in co-culture experiments using human monocyte-derived macrophages (MDMs) isolated from peripheral blood (Supplementary Figures 4 and 5). Previously, we reported functional AR in macrophages^37^. To ensure that our observations were truly intrinsic to the tumor cells and not influenced by the macrophages *in trans*, and to determine whether the mode of killing required cell-cell interactions or was caused by factors secreted by macrophages, we collected conditioned media (CM) of macrophages for cell viability assays. LNCaP cell viability was greatly reduced when cultured in androgen-proficient CM compared to androgen-deficient CM, suggesting that factors secreted by macrophages causes this effect and confirming that AR-action in tumor cells is responsible for susceptibility to macrophage-mediated killing and not AR in macrophages (Figure 5D). In addition, the co-culture results were successfully validated in a second AR-driven PCa cell line, LuCaP 35, established from the LuCaP 35 PDX model^39^ (Figure 5E). Taken together, these results indicate that AR acts as an immunomodulator, and that AR activity status determines whether cells are susceptible to macrophage-mediated cell death.

**Figure 5.**
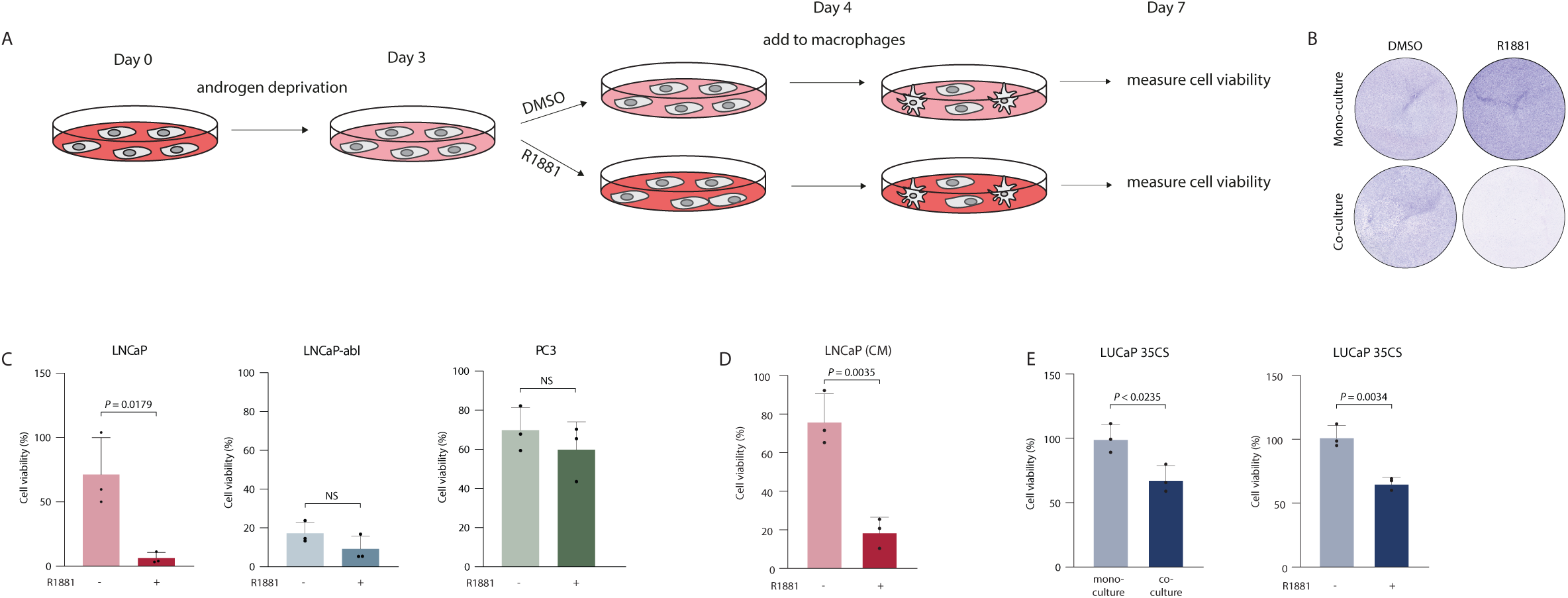
AR acts as an immunomodulator and regulates macrophage-mediated PCa cell killing. A. Schematic illustration of hormonal perturbation of LNCaP cells. Cells were deprived of androgens for three days by culturing in RPMI + 5% DCC + 1% Pen/Strep. After 3 days, cells were stimulated with either vehicle (DMSO) or 10nM R1881 for 24h (day 4). After 24h, cells were co-cultured with macrophages for 3 days and cell viability was measured using cell titer glo (day 7). B. Crystal violet staining of LNCaP cells cultured as a monoculture (top) or co-cultured with macrophages (bottom) pretreated with either vehicle (DSMO) or 100pM R1881. Prior to the start of the experiment, the cells were androgen deprived for 3 days. Subsequently, the same number of LNCaP cells were seeded in the monoculture condition (top) as in the co-culture condition (bottom). Cells were stimulated with either vehicle (DMSO) or 100pM R1881. Cells were stained with crystal violet after three days. C. Cell viability assay of LNCaP (left), LNCaP-abl (middle) and PC3 (right) cells pretreated with either vehicle (DMSO) or 100pM R1881 for 24h. Thereafter, cells were co-cultured with macrophages for 3 days after which cell viability was measured (n=3, mean values are depicted and error bars represent the standard deviation). Data was normalized to the monoculture condition. P-value was calculated using a two-tailed unpaired t-test. D. Cell viability assay of LNCaP cells pretreated with either vehicle (DMSO) or 100pM R1881 for 24h and subsequently cultured in conditioned medium (CM) of macrophages. Data is normalized to the normal medium condition (n=3, mean values are depicted and error bars represent the standard deviation). P-value was calculated using a two-tailed unpaired t-test. E. Cell viability assay of LUCaP35 cells. Left: cells cultured alone (monoculture) or in a co-culture with macrophages. Data is normalized to the monoculture condition (n=3, mean values are depicted and error bars represent the standard deviation). Right: LUCaP 35 cells pre-treated with either vehicle (DMSO) or 100pM R1881 for 24h. Thereafter, cells were co-cultured with macrophages for 3 days after which cell viability was measured (n=3, mean values are depicted and error bars represent the standard deviation). Data is normalized to the monoculture condition. P-value was calculated using a two-tailed unpaired t-test.

### Genes mediating ferroptosis resistance are upregulated by AR inhibition

Phospho-proteomic analyses indicated that AR stimulation activated the ferroptosis signaling pathway (Figure 4E), functionally connecting all genes identified in the screen. Indeed, AR activation by R1881 significantly decreased cell viability of LNCaP cells when treated with Erastin (VDAC2/VDAC3 inhibitor^40^), suggesting that androgens sensitize cells to ferroptosis-regulated cell death (Figure 6A).

**Figure 6.**
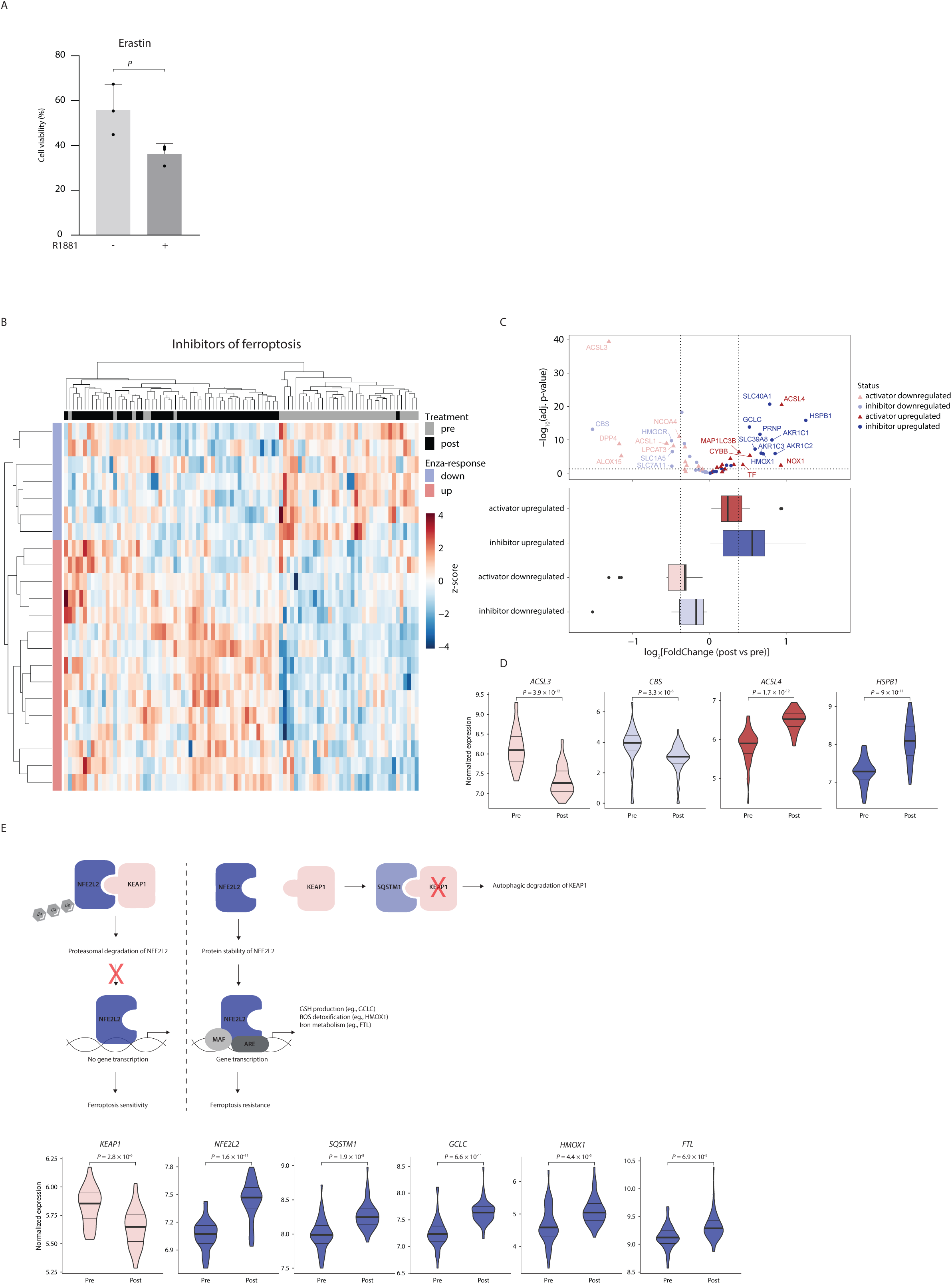
Genes mediating ferroptosis resistance are upregulated by AR inhibition. A. Cell viability assay of LNCaP cells pre-treated with either vehicle (DMSO) or 100pM R1881 for 24h. Thereafter, cells were treated with either vehicle (DSMO) or 10nM Erastin. Cell viability was measured after 3 days. Data is normalized to vehicle control (n=3, mean values are depicted and error bars represent the standard deviation). B. Unsupervised hierarchical clustering of pre-and post-treatment RNA-seq samples based on the expression of ferroptosis inhibitor genes obtained from the Humane Gene Set Ferroptosis (M39768, GSEA). Color scale indicates gene expression (z-score). C. Differential gene expression analysis of ferroptosis genes in prostate tumor samples post vs pre-enzalutamide treatment, in a volcano-plot (top) and a boxplot (bottom). Dark red: ferroptosis activator genes upregulated. Light red: ferroptosis activator genes downregulated. Dark blue: ferroptosis inhibitor genes upregulated. Light blue: ferroptosis inhibitor genes downregulated. Cut-off is depicted with a dotted line (|Fold Change| ≥ 1.3, p-adjusted < 0.05). D. Violin plots showing normalized gene expression of ACSL3 (activator downregulated), CBS (inhibitor downregulated), ACSL4 (activator upregulated) and HSPB1 (inhibitor upregulated) before (pre) and after 3 months (post) of neoadjuvant enzalutamide treatment. P-value was calculated using a Mann-Whitney U test. E. Schematic overview of the NFE2L2/KEAP1 pathway. Left: under normal conditions, low levels of NFE2L2 are maintained primarily by KEAP1-mediated proteasomal degradation. Right: following ferroptosis stress, NFE2L2 protein is stabilized and initiates a multi-step activation pathway, including nuclear translocation, heterodimerization with its partner MAF protein, recruitment of transcriptional coactivators, and subsequent binding to the antioxidant response element (ARE) of the target gene promoter. *SQSTM1* can stabilize *NFE2L2* by inactivating KEAP1 through autophagic degradation. Violin plots showing normalized gene expression of KEAP1 (activator downregulated), *NFE2L2* (inhibitor activated), *SQSTM1* (inhibitor activated), GCLC (inhibitor activated), *HMOX1* (inhibitor activated), *FTL* (inhibitor activated). P-value was calculated using a Mann-Whitney *U* test.

To confirm these observations in clinical samples, we next analyzed transcriptomics data from a phase II neoadjuvant clinical trial (DARANA, NCT03297385) enrolling high risk patients with localized PCa, where PCa tissue samples were taken before and after three months of monotherapy with the AR-inhibitor Enzalutamide^41^. Genes from the ferroptosis gene set (Human Gene Set; #M39768) were divided into ferroptosis activators and inhibitors, based on literature (Supplementary Figure 6). Interestingly, the majority of ferroptosis inhibitor genes were upregulated upon enzalutamide treatment (Figure 6B). Furthermore, when we analyzed the expression of the ferroptosis regulator genes after enzalutamide treatment, we found that the majority of ferroptosis inhibitor genes were upregulated after enzalutamide treatment, while the other subclasses did not change significantly (Figure 6C and D). These results were not observed for the ferroptosis activator genes (Supplementary Figure 7).

NFE2L2/KEAP1 signaling plays a critical role in ferroptosis sensitivity and resistance, as nearly all genes thus far implicated in ferroptosis are transcriptionally regulated by NFE2L2^42, 43^ (schematic overview in Figure 6E). NFE2L2 is a transcription factor that coordinates the activation of cell protection genes involved in iron metabolism, redox signaling and oxidative defense during ferroptosis^44^. Under normal conditions, the kelch-like ECH-associated protein 1 (KEAP1) ubiquitinates NFE2L2 and mediates its proteasomal degradation, rendering the cell sensitive to ferroptosis^42^. However, following ferroptosis stress, NFE2L2 is stabilized by *SQSTM1*, whereupon NFE2L2 heterodimerizes with its interaction partner MAF to drive genes involved in GSH production, ROS detoxification and iron metabolism, all of which result in ferroptosis resistance (Figure 6E). Therefore, we evaluated this signaling pathway in our clinical trial samples, comparing pre-and post-enzalutamide treatment specimens. Strikingly, expression of the ferroptosis inhibitor KEAP1 was significantly downregulated after enzalutamide treatment, and expression of both *NFE2L2* and its stabilizer *SQSTM1*, mediating ferroptosis resistance^42^, were significantly upregulated after enzalutamide treatment. In addition, the expression of NFE2L2 target genes *GCLC, HMOX1* and *FTL* were all significantly upregulated upon enzalutamide treatment. This indicates that AR-targeted therapy activates genes critical for ferroptosis resistance, ultimately protecting the tumor cell from macrophage-mediated killing capacity. In summary, we observed that AR-activation status is inversely associated with ferroptosis resistance, both in cell lines and in patients. These data imply that AR-targeted therapy - the mainstay treatment for PCa patients - has an unexpected and previously unknown adverse effect: it prevents tumor cells from being killed by macrophages, through ferroptosis induction.

## Discussion

Treatment with ADT and anti-androgens is the mainstay therapy for patients with locally advanced, metastatic and biochemically recurrent PCa. Unfortunately, resistance to therapy often occurs leading to the emergence of CRPC, which is inevitably lethal. Therefore, the discovery of new treatment regimens is vital to increase the overall survival of patients with advanced PCa.

The tumor microenvironment plays a pivotal role in tumor development and progression and may provide novel therapeutic opportunities. Specifically, the interaction between PCa cells and macrophages is critically involved in the growth and spread of PCa^45, 46^, and represents a promising therapeutic avenue^47, 48^. For example, inhibitors of the colony-stimulating factor 1 receptor (CSF1R), which is involved in macrophage recruitment and polarization, reduce tumor growth in preclinical models of PCa^49–51^. In addition, therapeutics that modulate macrophage-cancer cell interactions, have shown clinical efficacy^52, 53^. Here, we identified that AR activity in the tumor cells is a key requirement for macrophage-mediated tumor killing capacity. This observation may have far- reaching consequences and highlights a direct adverse effect of hormonal targeting in PCa, which diminishes the anti-tumor potency of the innate immune system.

Recently, bipolar androgen therapy (BAT) has been introduced in clinical trials with promising results^52, 53^, but the underlying mechanism remains incompletely understood. Our results may provide a plausible mechanism of action for this therapeutic strategy, in which the tumor growth reducing effect of ADT (but preventing tumor killing by macrophages) is alternated with the tumor-killing effect of AR-activity (but stimulating tumor growth). Although plausible, future studies should be designed to formally test this hypothesis.

Pro-inflammatory macrophages can support tumor-cell killing in three different ways: (1) indirectly by the recruitment of other immune cells, (2) through antibody-dependent cellular cytotoxicity (cell contact) and (3) direct killing through the secretion of harmful products (e.g., cytokines and oxygen radicals)^40–42^. Since our study shows that conditioned medium from macrophages is sufficient to kill tumor cells, being disrupted when the tumor cells are deprived of hormone, we conclude that factors secreted by macrophages represent the critical mediator in the observed response. Our results show that NF-κB components are key regulators of macrophage-mediated killing in PCa cells. Indeed, factors secreted by macrophages include compounds that activate NF-κB and PKCδ signaling, including TNFα, IL-1, and reactive oxygen species (ROS)^54–56^. TNFα induces PKCδ-mediated apoptosis in androgen-dependent PCa cells, through the JNK and p38 mitogen-activated protein kinase cascades^57–59^. In addition, AR inhibits NF-κB responses^60–62^. Interestingly, elevated levels of TNFα in PCa correlate with poor prognosis^63^ and the NF-κB pathway may also promote resistance to androgen deprivation^64, 65^. Amplification of AR and activation of the NF-κB pathway are often observed in CRPC, whereas in this study, we demonstrated that inhibition of AR signaling or reduced NF-κB signaling results in PCa cells that are resistant to macrophage-mediated killing, implying that AR and NF-κB signaling play different roles depending on the context, whether it is intrinsic to the tumor cell or dependent on the tumor microenvironment.

Here, we showed that the ferroptosis signaling pathway is downregulated upon perturbation of NF-κB and PRKCD signaling, conferring resistance to macrophage-mediated killing. Interestingly, there is a crosstalk between NF-κB signaling and the NRF2 pathway, which regulates the oxidative-stress response and important in ferroptosis resistance^66^. In addition, phosphorylation of NRF2 at serine 40 by PRKCD facilitates NRF2 nuclear translocation and ferroptosis sensitivity^67^. These findings highlight that both NFκB and PKCδ are critically involved in the ferroptosis signaling pathway and that disruption of these proteins confers ferroptosis resistance.

In the past years, the regulatory relationship of AR in ferroptosis signaling has gained scientific interest. AR increases the expression of STAMP2, driving the reduction of Fe^3+^ to Fe^2+^, while depleting available NADPH, leading to increased intracellular ROS levels and induced ferroptosis^68^. In addition, androgens regulate the expression of SREBP1, the master regulator of lipid metabolism which controls SCD1, a master regulator of ferroptosis^69, 70^. Our findings link AR signaling to macrophage-mediated killing capacity, through ferroptosis. While the connection of macrophage-mediated killing and AR action is novel, a functional connection between AR and ferroptosis has recently been reported by others via MBOAT2; an androgen-responsive gene^71^. However, as our genome-wide CRISPR-screen did not identify MBOAT2 as a hit, functionally redundant cascades may exist in parallel, that are jointly under the control of AR as a master regulator.

Our studies identified AR activation as essential for macrophage-mediated tumor cell death, establishing AR as a true immune modulator and highlighting immune protection from macrophages as a negative side effect of hormone therapy in PCa patients. The constant interactions between tumor cells and the microenvironment play a critical role in tumor initiation, progression, metastasis and response to therapy. Future studies should be geared to restore macrophage-mediated tumor killing capacity despite hormonal intervention, enhancing therapeutic efficacy and preventing disease relapse, a common feature currently observed for practically all prostate cancer patients receiving AR-targeted therapies.

## Acknowledgements

We thank members of the Zwart and Bergman labs for suggestions, input and valuable feedback. We would like to thank the NKI Genomics Core Facility for next-generation sequencing and bioinformatics support. We thank the NKI Proteomics/Mass Spectrometry Facility. We thank Reuven Agami, Julien Champagne and Remco Nagel for fruitful discussions and technical support. We thank Peter Nelson for technical support (P.S.N. was supported by research grants P50CA097186, P01CA163227 and R01CA266452)

## Author contributions

Conceptualization: A.Z., W.Z., and A.M.B.; W.Z. and A.M.B. were responsible for project funding; A.Z. I.M.P., and K.S. performed the CRISPR-screen; C.L. and R.L.B. provided bioinformatics and technical support for the CRISPR screen; L.H. and M.A. performed and analyzed the proteomic experiments; A.Z. and T.A.v.S. performed co-culture experiments; A.Z performed cell proliferation, crystal violet and luciferase experiments; A.Z performed MDM isolation; A.Z. performed CM assays; A.Z. and E.M. performed RT-qPCR and western blot analyses. E.M. and S.G. performed CUT&Tag experiments. A.Z. and S.G. performed flow cytometry experiments and analysis. A.Z., W.Z., and A.M.B. wrote the manuscript, with input from all co-authors.

## Materials and methods

### Cell lines

The prostate cancer cell lines LNCaP and PC3 and the monocytic cell line THP-1 were cultured in Roswell Park Memorial Institute (RPMI) 1640 medium supplemented with 10% fetal bovine serum (FBS) and 1% penicillin-streptomycin (Pen/Strep) (5000 U/mL, life technologies). The LNCaP derived castration resistant clone, LNCaP-abl (ablated) was cultured in RPMI medium supplemented with 5% dextran coated charcoal (DCC) stripped serum and 1% Pen/Strep. These cell lines were purchased from American Type Culture Collection (ATCC). The LuCaP 35CS cell line (kindly provided by Peter Nelson, Fred Hutchinson Cancer Center), derived from the LuCaP35 patient derived xenograft, was cultured in Dulbecco’s Modified Eagle Medium (DMEM) supplemented with 10% FBS and 1% Pen/Strep. For hormonal related experiments, all cells were cultured 3 days prior to the start of the experiment in medium supplemented with 5% DCC and 1% Pen/Strep. AR was induced with 10nM R1881 (synthetic androgen, Sigma) unless otherwise stated. All cell lines were cultured at 5% CO_2_ and 37 °C. All cell lines were genotyped and routinely tested for mycoplasma infection.

### Macrophage differentiation

THP-1 cells were stimulated with 100ng/mL of phorbol 12-myristate 13-acetate (PMA, Sigma) for 48h, followed by 24h in fresh RPMI supplemented with 10%FBS and 1%Pen/Strep. After 24h, cells were differentiated by a 24h stimulation of 10ng/mL lipopolysaccharide (LPS, Sigma) and 10ng/mL interferon-γ (IFN-γ, Peprotech).

For AR signaling experiments, THP-1 cells were stimulated with 100ng/mL of phorbol 12-myristate 13-acetate for 48h, followed by 24h in fresh RPMI supplemented with 5%DCC and 1%Pen/Strep. After 24h, cells were differentiated by a 24h stimulation of 10ng/mL lipopolysaccharide and 10ng/mL interferon-γ. Thereafter, cells were gently washed with PBS and stimulated with vehicle or 10nM R1881 for 24h.

### Conditioned medium generation and collection

For the generation of conditioned medium (CM), THP-1 cells were stimulated with 100ng/mL of phorbol 12-myristate 13-acetate for 48h, followed by 24h in fresh RPMI supplemented with 10%FBS and 1%Pen/Strep. After 24h, cells were differentiated by a 24h stimulation of 10ng/mL lipopolysaccharide and 10ng/mL interferon-γ. After 24h, cells were gently washed with PBS and fresh medium was added. CM was harvested after 48h, centrifuged, and filtered through a 0.45 μm strainer. CM was either freshly used or snap frozen in liquid nitrogen and stored at −80 °C.

For AR signaling experiments, THP-1 cells were stimulated with 100ng/mL of phorbol 12-myristate 13-acetate for 48h, followed by 24h in fresh RPMI supplemented with 5%DCC and 1%Pen/Strep. After 24h, cells were differentiated by a 24h stimulation of 10ng/mL lipopolysaccharide and 10ng/mL interferon-γ. Thereafter, cells were gently washed with PBS and stimulated with vehicle or 10nM R1881 for 24h. After 24h, cells were gently washed with PBS and fresh medium was added. CM for vehicle and R1881-stimulated conditions was harvested after 48h, centrifuged, and filtered through a 0.45 μm strainer. CM was either freshly used or snap frozen in liquid nitrogen and stored at −80 °C.

### Generation of monocyte-derived macrophages

Whole peripheral blood was obtained from healthy male donors. Buffy coat was mixed 1:2 with PBS and added onto a Ficoll gradient in a 3:1 ratio (Invitrogen). This was centrifuged at 2100 rpm for 25 minutes at RT (w/o brakes). The leukocyte ring was collected and washed with cold PBS. Cells were then resuspended in 0.5%BSA-PBS containing CD14 microbeads (Miltenybiotec). CD14+ cells were cultured in RPMI medium supplemented with 10% FBS, 1% Pen/Strep and 20ng/mL GM-CSF (Miltenybiotec) for 96h. Thereafter, cells were stimulated with 10ng/mL lipopolysaccharide and 10ng/mL interferon-γ for differentiation into pro-inflammatory macrophages. Cells were subsequently checked for pro-inflammatory markers by flow cytometry (Supplementary Figure 4).

For AR signaling experiments, CD14^+^ cells were isolated as described above. Subsequently, CD14+ cells were cultured in RPMI medium supplemented with 10% FBS, 1% Pen/Strep and 20ng/mL GM-CSF (Miltenybiotec) for 24h. Thereafter, medium was replaced with RPMI supplemented with 5% DCC, 1% Pen/Strep and 20ng/mL GM-CSF for 3 days. Cells were then stimulated with 10ng/mL lipopolysaccharide and 10ng/mL interferon-γ for differentiation into pro-inflammatory macrophages. Thereafter, cells were stimulated with vehicle or 10nM R1881 for 24h. After 24h, cells were used for co-culture experiments.

### Genome wide co-culture CRISPR-screen

The human genome wide CRISPR library Brunello^15, 16^ was used, which was a kind gift from Roderick Beijersbergen (The Netherlands Cancer Institute, NKI). The lentiviral Brunello library was transduced into low passage LNCaP cells at a low multiplicity of transduction (MOI) of ∼0.3 to ensure that only one sgRNA was incorporated per cell. After 48h, cells were selected with 2ug/mL puromycin for 72h and left untreated for 10 days. Thereafter cells were either used as reference population (T0) or cultured for 72h with macrophages (co-culture). For T0, 32×10^6^/mL cells/arm were isolated to ensure a 400x coverage of the library. For the co-culture, all cells after 72h were isolated. Genomic DNA was subsequently isolated according to manufacturer’s instructions using the Gentra Puragene Cell kit. gRNAs were amplified by two consecutive PCRs as previously described (Korkmaz et al., 2016). DNA libraries were sequenced on HiSeq 2500 platform (single-read; 65bp). sgRNA abundance was analyzed using DESeq2 and MAGeCK tools Robust Rank Algorithm (RRA)^72, 73^.

For the T0 control arm, regarding the distribution of the sgRNA counts within a sample, almost all sgRNAs have at least 10 counts and approximately 62% of the sgRNAs have a count above 100 (Supplementary Figure 1C). In the tumor cell/macrophage co-culture arm, approximately 50% of all sgRNAs were depleted, with a count of less than 10 (Supplementary Figure 1C).

For sequence depth normalization, a relative total size factor was calculated for each sample, by dividing the total counts of each sample by the geometric mean of all totals as described^74^. After normalization, a differential test between the treated (co-culture) and untreated (T0) condition for each sgRNA was performed using DESeq2, and the output was sorted. MAGeCK RRA was used to determine for each gene whether its sgRNAs were enriched toward the top of the result list. The resulting enrichment P values were corrected for multiple testing using the Benjamini-Hochberg correction, resulting in a false-discovery rate (FDR)-corrected value. Hits were selected based on the following criterion: the comparison between co-culture vs T0: FDR £ 0.1 and the |FcMedian| ³ 1.5. Results of the screen can be found in Supplementary Table 1.

### Generation of CRISPR knockout cell lines

To generate CRISPR knockout cell lines, sgRNAs were selected from the Brunello and the GeCKOv2 human CRISPR KO library to target CHUK (sgRNA_1: CACCGACAGACGTTCCCGAAGCCGC, sgRNA_2: CACCGTTCTGGAGGAGATCTCCGAA), IKBKB (sgRNA_1: CACCGGCCATGGAGTACTGCCAAGG, sgRNA_2: CACCGTCAGCCCCCGGAACCGAGAG), IKBKG (sgRNA_1: CACCGGAGGAGAATCAAGAGCTCCG, sgRNA_2: CACCGTCAGGAGCGCCCTGTTCTGA), PRKCD (sgRNA_1: CACCGTGCAGAGCGTGGGAAAACAC, sgRNA_2: CACCGTTCCCAACGATGAACCGCCG), and non-targeting (NT) (sgRNA_1: CACCGAAATGCACAGATCGCTGATC, sgRNA_2: CACCGAACGCTGTCGTACGTGTATA, sgRNA_3: CACCGAACTACAAGTAAAAGTATCG). These sgRNAs were cloned into the lentiCRISPRv2 plasmid as previously described^75^. After cloning, all constructs were verified by Sanger sequencing. Lentivirus was generated in HEK293T cells cultured in DMEM supplemented with 10% FBS and 1% Pen/Strep.

HEK293T cells were transfected using polyethylenimine (PEI, 1mg/mL) with three viral packaging constructs (pRC/CMV-rev1B, pHDM-Hgpm2 and pHDM-G) and lentiCRISPRv2 constructs. 3.5×10^6^ HEK293T cells were seeded in 10cm^2^ plates and incubated overnight. After 24h, viral vectors were produced by mixing the three packaging constructs in a 1:1:1 ratio, and 10.5 μg of packaging mix was added to 17.5 μg of the respective lentiCRISPRv2 construct with 105 μl of PEI, incubated for 8 mins and added to the HEK293T cells. After overnight incubation, cells were refreshed with 8mL of medium. After 24h, the supernatant was harvested, filtered with a 0.22 μm filter (Millipore) and added onto low passage LNCaP cells with polybrene (8μg/mL). After 72hrs, cells were selected with puromycin (2μg/mL).

### PRKCD Hi-ChIP and CUT&Tag

H3K27ac HiChIP sequencing regions from LNCaP cells^33, 34^ were downloaded and filtered for promoter-enhancer interactions. Filtered anchors were intersected with AR binding sites found in LNCaP cells^32^ (GSM2480801). The CUT&Tag experiment was performed according to the CUT&Tag Protocol (v1.7) as previously described by EpiCypher (derived from Kaya-Okur et al.^76^). In brief, we isolated 100,000 unfixed nuclei from LNCaP cells per condition which were incubated with, and thereby adsorbed to, activated Concanavalin A beads. After adsorption, the bead/nuclei mixture was incubated overnight with a 1:100 dilution of primary antibody (H3K27ac (39133, Active Motif)) which was followed by secondary antibody (13-1047, EpiCypher) incubation. Next, the pAG-Tn5 was added to each reaction which, together with the tagmentation buffer, lead to the tagmentation of the DNA. After DNA amplification (15-cycles) and library preparation, the samples were sequenced on the Illumina NextSeq 550 platform using the paired-end protocol (44bp * 32 bp) and aligned to the human reference genome hg38.

### RNA isolation, reverse-transcription and quantitative real-time PCR (RT-qPCR)

Cells were gently washed with PBS before total RNA was isolated using Invitrogen^TM^ TRIzol^TM^ Reagent (Thermo Fisher Scientific) according to the manufacturer’s protocol. First-strand cDNA was synthesized from 2 μg RNA using SuperScript^®^ III First-Strand Synthesis System for Reverse Transcriptase-PCR (Life Technologies) with random hexamer primers according to the manufacturer’s instructions. RT-qPCR was performed using SensiMix™ SYBR® No-ROX Kit (Bioline) in a QuantStudio™ 6 Flex System (Thermo Fisher Scientific). A list of primers can be found in Supplementary Table 2.

### Western blot

Cells were gently washed with PBS before protein was isolated using 2x Laemmli buffer (120 mM Tris, 20% glycerol, 4% SDS) supplemented with protease inhibitor (1:100) and phenylmethylsulfonyl fluoride (PMSF, 1:200). Lysates were sonicated (EpiShear Probe Sonicatore, Active Motif) for 10 cycles with one second intervals and a 20% amplitude using. Per sample, 30μg of protein was resolved by SDS-PAGE (10%) in SDS-PAGE 1X Running buffer (25 mM Tris, 0.25 M glycine, 0.1% SDS) and transferred on nitrocellulose membranes (Santa Cruz Biotechnology) in cold 1X Transfer buffer (24 mM Tris, 192 mM glycine) on ice at 100 V for 90 min or at 0.9 mA overnight at 4°C . After transfer, membranes were stained with Ponceau S and subsequently blocked in 4% BSA (A8022, Sigma/Merck) in 1x PBS-Tween (137 mM NACl, 10 mM Na_2_HP04, 1.5 mM KH_2_PO_4_, 2.6 mM KCl, 0.1% Tween-20) for 1h and incubated with primary antibodies against IKKα (2682, Cell signaling, 1:1000), IKKβ (10AG2, Novus Biological, 1:000), IKKγ (2685, Cell signaling, 1:1000), PKCδ (2058, Cell signaling, 1:1000), Actin (MAB1501R, Merck, 1:1000), HSP90 (sc-12119, Santa Cruz Biotechnologies, 1:1000) diluted in 4% BSA/PBS-T. Blots were incubated overnight at 4°C and washed five times with PBS-T prior to incubation with secondary antibodies donkey-α-mouse 680 RD (926-68073, LI-COR Biosciences, 1:10000) and donkey-α-rabbit 800 CW (926-32213, LI-COR Biosciences, 1:10000), diluted in 4% BSA/PBS-T for 1h. Membranes were scanned and analyzed using and Odyssey^®^ CLx Imaging System (LI-COR Biosciences) and ImageStudio^™^ Lite v5.2.5 software (LI-COR Biosciences).

### In vitro experiments

#### Cell proliferation assay

LNCaP cells were seeded in a 384-well plate (CELLSTAR plate, 384w, F, vClear, TC, PS, black, lid, Greiner) at a density of 3000 cells/well. For co-culture experiments, THP-1 cells were seeded in a 1:4 ratio (750 cells/well) and differentiated into pro-inflammatory macrophages as described above. Cells were imaged every 4h using an IncuCyte Zoom Live-Cell Analysis System. Cell confluency percentage was calculated using the IncuCyte Zoom software.

#### Crystal violet assay

Cells were seeded in a 6-well plate at a density of 500,000 cells/well. After three days of co-culture, cells were fixed using ice-cold MeOH for 10 minutes. Subsequently, cells were gently washed with PBS and stained with 0.1% crystal violet for 1hr. Thereafter, cells were gently washed with PBS and dried at RT prior to scanning.

#### Co-culture experiments

For co-culture experiments, THP-1 cells were differentiated into pro-inflammatory macrophages as described above. Upon differentiation, LNCaP, LNCaP-abl, PC3 or LuCaP cells were added to the macrophages in a 1:4 (E:T) ratio.

For AR signaling experiments, PCa cells were hormone deprived for 3 days by culturing the cells in RPMI + 5% DCC + 1% Pen/Strep. Thereafter, cells were stimulated with vehicle (DMSO) or 100pM of R1881 for 24h and added to the macrophages.

#### Cell viability assay

For cell viability assays, LNCaP, LNCaP-abl, PC3 or LuCaP35 cells were seeded at 3000 cells/well in a 384-well plate (CELLSTAR plate, 384w, F, vClear, TC, PS, black, lid, Greiner). For co-culture experiments, THP-1 cells were seeded in a 1:4 ratio (750 cells/well) and differentiated into pro-inflammatory macrophages as described above. For co-culture experiments, cell viability was assessed 3 days after the start of the co-culture. For Erastin experiments, LNCaP cells were hormone deprived for 3 days by culturing the cells in RPMI + 5% DCC + 1% Pen/Strep. Thereafter, cells were stimulated with vehicle (DMSO) or 100pM of R1881 for 24h. After 24h, cells were stimulated with 10μM of Erastin. Cell viability was assessed 5 days after the start of the treatment. Cell viability was assessed using the CellTiter-Glo Luminescent Cell Viability Assay kit (Promega) per the manufacturer’s instructions. Bar charts were plotted using GraphPad Prism 9 software.

#### Reporter gene assay for NF-κB

To investigate NF-κB signaling activity in the LNCaP KO and NT cells, a luciferase reporter gene assay was used. LNCaP KO and LNCaP NT control cells were transiently transfected with 150ng of NF-kB luciferase reporter gene (NF-κB-luc, Clontech) and 3.5ng of Renilla using Lipofectamine 2000 (Invitrogen) according to the manufacturer’s instructions in a 96-well plate (Greiner) at a density of 15,000 cells per well for 24h. Cells were stimulated with either vehicle (H_2_O) or 15ng/mL TNF-1 or normal medium (NM) and CM for 24h. After 24h, cells were lysed using the Dual-Luciferase^®^

Reporter Assay System (Promega) and luciferase activity was measured using a Tecan plate reader (Infinite M Plex. Tecan).

#### Flow Cytometry

MDMs were differentiated into pro-inflammatory macrophages as described above. After differentiation, cells were harvested by scraping. Subsequently, cells were washed with PBS, resuspended in 2% Bovine Serum Albumin (BSA)-PBS, strained through a 0.35μm cell strainer and placed on ice. Next, cells were labeled with an antibody cocktail containing Anti-CD14-PE/Cy7 (Clone 63D3, 367111, BioLegend, 1:100), Anti-CD80-APC (Clone 2D10, 305219, BioLegend, 1:100), Anti-CD86-FITC (Clone BU63, 374203, BioLegend, 1:100) for 45 minutes on ice. Cells were washed with PBS and resuspended in 2%BSA-PBS and checked on a Fortessa flow cytometer. DAPI was added just before running the sample to check cell viability. Flow cytometry analysis was performed using FlowJo^TM^ Software (BD Biosciences).

#### (Phospho)proteomic analyses

For protein digestion, frozen cell pellets were lysed in boiling Guanidine (GuHCl) lysis buffer as previously described^77^. Protein concentration was determined with a Pierce Coomassie (Bradford) Protein Assay Kit (Thermo Scientific), according to the manufacturer’s instructions. Aliquots corresponding to 1.1mg of protein were digested with Lys-C (Wako) for 21h at 371°C, enzyme/substrate ratio 1:100. The mixture was then diluted to 2M GuHCl and digested overnight at 37 °C with trypsin (Sigma-Aldrich) in enzyme/substrate ratio 1:100. Digestion was quenched by the addition of TFA (final concentration 1%), after which the peptides were desalted on a Sep-Pak C18 cartridge (Waters, Massachusetts, USA). From the eluates, aliquots were collected for proteome analysis, the remainder being reserved for phosphoproteome analysis. Samples were vacuum dried and stored at −801°C until LC-MS/MS analysis or phosphopeptide enrichment.

Phosphorylated peptides were enriched from 1mg of total peptides using High-Select Fe-NTA Phosphopeptide Enrichment Kit (Thermo Scientific), according to the manufacturer’s instructions, with the exception that the dried eluates were reconstituted in 15μl of 2% formic acid.

Prior to mass spectrometry analysis, the peptides used for proteome analysis were reconstituted in 2% formic acid. Peptide mixtures were analysed by nanoLC-MS/MS on an Q Exactive HF-X Hybrid Quadrupole-Orbitrap Mass Spectrometer equipped with an EASY-NLC 1200 system (Thermo Scientific). Samples were directly loaded onto the analytical column (ReproSil-Pur 120 C18-AQ, 1.9μm, 751μm1×15001mm, packed in-house). Solvent A was 0.1% formic acid/water and solvent B was 0.1% formic acid/80% acetonitrile. Samples were eluted from the analytical column at a constant flow of 2501nl/min. For single-run proteome analysis, a 4-h gradient was employed containing a linear increase from 7 to 30% solvent B, followed by a 15 min wash, whereas for single-run phosphoproteome analysis, a 2-h linear gradient (from 4 to 22% solvent B, followed by a 15-min wash) was used.

Proteome data was analysed by PD (v. 2.3.0.523, Thermo Scientific) using standard settings. MS/MS data were searched against the human Swissprot database (20417 entries, release 2019_02) using Sequest HT. The maximum allowed precursor mass tolerance was 50 ppm and 0.061Da for fragment ion masses. Trypsin was chosen as cleavage specificity allowing two missed cleavages. Carbamidomethylation (C) was set as a fixed modification, while oxidation (M) and deamidation (NQ) were used as variable modifications. False discovery rates for peptide and protein identification were set to 1%, and as an additional filter Sequest Ht XCorr>1 was set. The PD output file containing the abundances was loaded into Perseus (v. 1.6.1.3) [02]. LFQ intensities were Log2-transformed and the proteins were filtered for at least two out of three valid values in one condition. Missing values were replaced by imputation based on the standard settings of Perseus, i.e., a normal distribution using a width of 0.3 and a downshift of 1.8. Differentially expressed proteins were determined using a t test (threshold: P1≤10.05 and [x/y] ≥1.51|1[x/y] ≤−1.5).

Phosphoproteome data were analysed by MaxQuant (v. 1.6.1.0) using standard settings^78^. MS/MS data were searched against the human Swissprot database (20,417 entries, release 2019_02) complemented with a list of common contaminants and concatenated with the reversed version of all sequences. The maximum allowed mass tolerance was 4.5 ppm in the main search and 20 ppm for fragment ion masses. False discovery rates for peptide and protein identification were set to 1%. Trypsin/P was chosen as cleavage specificity allowing two missed cleavages. Carbamidomethylation (C) was set as a fixed modification, while oxidation (M), deamidation (NQ) and phosphorylation (S,T,Y) were used as variable modifications. LFQ intensities were Log2-transformed in Perseus (v. 1.6.5.0), after which the phosphosites were filtered for at least two valid values (out of 3 total) in both conditions. Missing values were replaced by imputation based on a normal distribution using a width of 0.3 and a downshift of 1.8. Differentially regulated phosphosites were determined using a ttest. These differential phosphosites were combined with on/off (three out of three total present/missing) phosphosites.

---

**Supplementary Figure 1. Quality control of the CRISPR-screen**

A. Representation of the library. Boxplot of the sgRNA counts in the Brunello library (77041) and in our data (T0, 76994). The number of sgRNAs are depicted on the y-axis.

B. Normalization of the sgRNA counts within the samples. A relative factor based on the total counts of each sample was calculated to normalize the values. The replicates are depicted on the x-axis and the total counts are depicted on the y-axis.

C. Distribution of the counts per sgRNA per sample. The distribution of sgRNAs of the three replicates individually of the T0 arm are depicted in red, the distribution of the sgRNAs of the three replicates individually of the co-culture (lm) arm are depicted in green. The rank is depicted on the x-axis and the log10Count is depicted on the y-axis.

**Supplementary Figure 2. IKK complex KO cells show reduced NF-κB luciferase activity.**

Luciferase reporter assay in NT cells and IKBKB.1-KO, IKBKB.2-KO, IKBKG.1-KO, IKBKG.2-KO, CHUK.1-KO and CHUK.2-KO cells. Cells were co-transfected with a NF-κB luciferase reporter and renilla. 24h after transfections, cells were stimulated with vehicle (H_2_O) or 15ng/mL TNFα (left) or with normal medium (NM) or conditioned medium (CM) for 24h. NF-κB activity was determined by luciferase assay. Normalized values (Luciferase over Renilla) are depicted on the y-axis (n=3, mean values are depicted and error bars represent the standard deviation). P-value was calculated using a two-way ANOVA test.

**Supplementary Figure 3. Cell proliferation is not affected in the KO cells.**

Cell viability assay of NT control cells and PRKCD.1-KO, IKBKB.1-KO, IKBKG.1-KO and CHUK.1-KO cells. Cell viability was measured after three days. RLU = relative light units. (n=3, mean values are depicted and error bars represent the standard deviation).

**Supplementary Figure 4. Isolation and differentiation of monocyte-derived-macrophages (MDMs).**

A. Workflow showing the generation of MDMs isolate from peripheral blood buffy coats of healthy donors.

B. Flow cytometry analysis of surface markers CD14, CD80 and CD86 at day 0 (up) and day 4 (down) of differentiation. Left: flow cytometry analysis of healthy donor (HD) 1. Right: flow cytometry analysis of HD2. Unstained cells (grey) were used as a negative control, HD samples are depicted in blue.

**Supplementary Figure 5. Monocyte-derived-macrophages can kill PCa cells depending on their AR status.**

A. Workflow showing the procedure of differentiation and AR stimulation of monocyte-derived-macrophages (MDMs). MDMs were stimulated with GM-CSF (day 0) in FBS containing medium. After 24h, cells were carefully washed with PBS and replenished with fresh DCC medium. Cells were stimulated with IFNγ and LPS (day 4) and vehicle (DMSO) or R1881 (day 5). After 24h of stimulation, cells were carefully washed and replenished with fresh medium and then co-cultured with different PCa cells.

B. Schematic illustration of hormonal perturbation of LNCaP, LNCaP-abl and PC3 cells. Cells were deprived of androgens for three days by culturing in RPMI + 5% DCC + 1% Pen/Strep. After 3 days, cells were stimulated with either vehicle (DMSO) or 10nM R1881 for 24h (day 4). After 24h, cells were co-cultured with macrophages for 3 days and cell viability was measured using cell titer glo (day 7).

C. Cell viability assay of LNCaP (left), LNCaP-abl (middle) and PC3 (right) cells pretreated with either vehicle (DMSO) or 100pM R1881 for 24h. Thereafter, cells were co-cultured with MDMs for 3 days after which cell viability was measured. Co-cultures with HD1 are shown at the top, co-cultures with HD2 are shown at the bottom (n=3, mean values are depicted and error bars represent the standard deviation). Data was normalized to the monoculture condition. P-value was calculated using a two-tailed unpaired t test.

**Supplementary Figure 6. Network of ferroptosis genes.**

String analysis (http://string-db.org) of the proteins that regulate ferroptosis (obtained from GSEA, Human Gene Set; #M39768). Red circles represent proteins that are known to activate ferroptosis, blue circles represent genes that are known to inhibit ferroptosis.

**Supplementary Figure 7. Heatmap of ferroptosis activating genes in prostate tumors pre-and post-enzalutamide treatment.**

Unsupervised hierarchical clustering of pre-and posttreatment RNA-seq samples based on the expression of ferroptosis activating genes obtained from the Humane Gene Set Ferroptosis (M39768, GSEA). Color scale indicates gene expression (z-score).

