## Supplementary figures and images for "A genome-wide CRISPR screen in human prostate cancer cells reveals drivers of macrophage-mediated cell killing and positions AR as a tumor-intrinsic immunomodulator"

### Supplementary Figure 1

#### Supplementary Figure 1

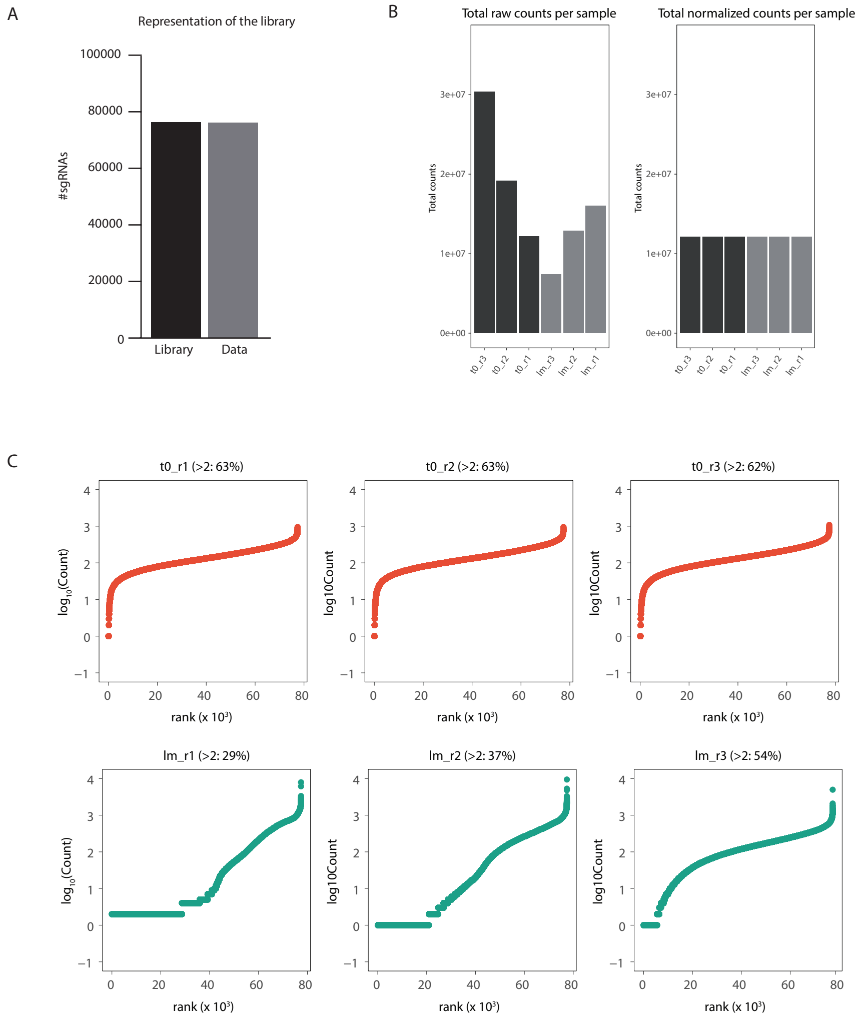

### Supplementary Figure 2

#### Supplementary Figure 2

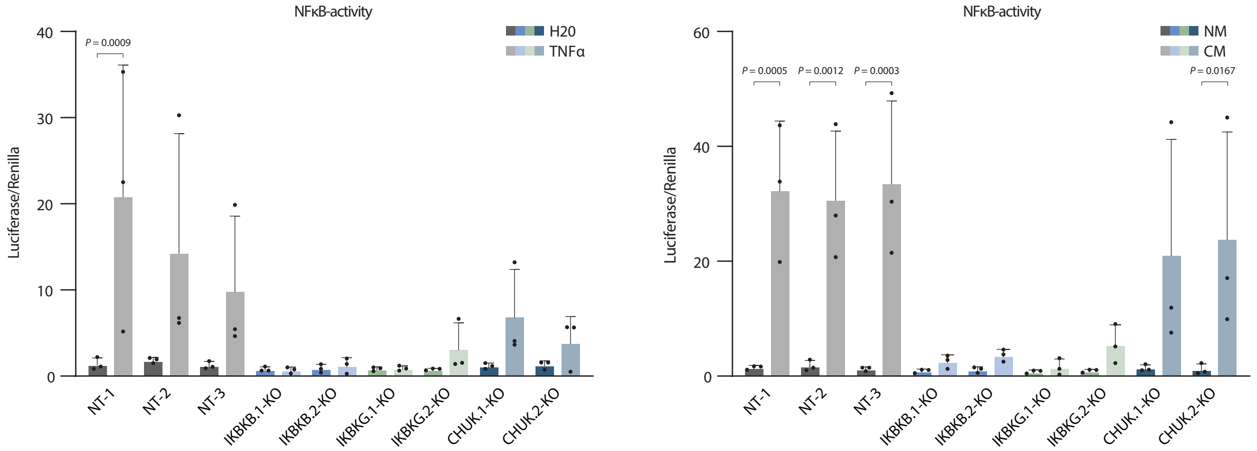

### Supplementary Figure 3

#### Supplementary Figure 3

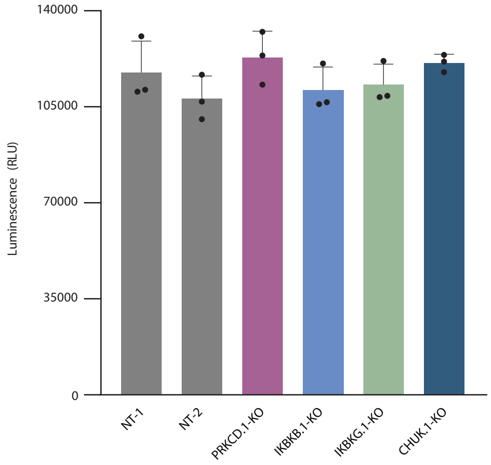

### Supplementary Figure 4

Supplementary Figure 4

A

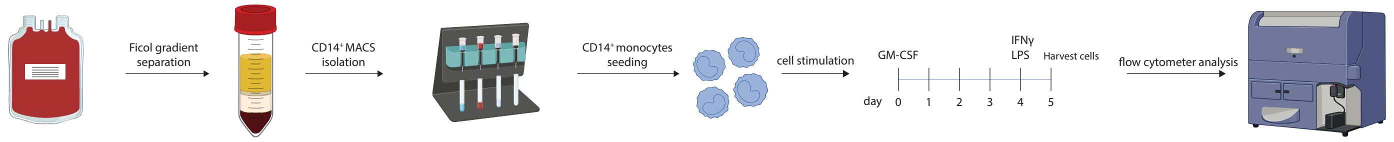


B

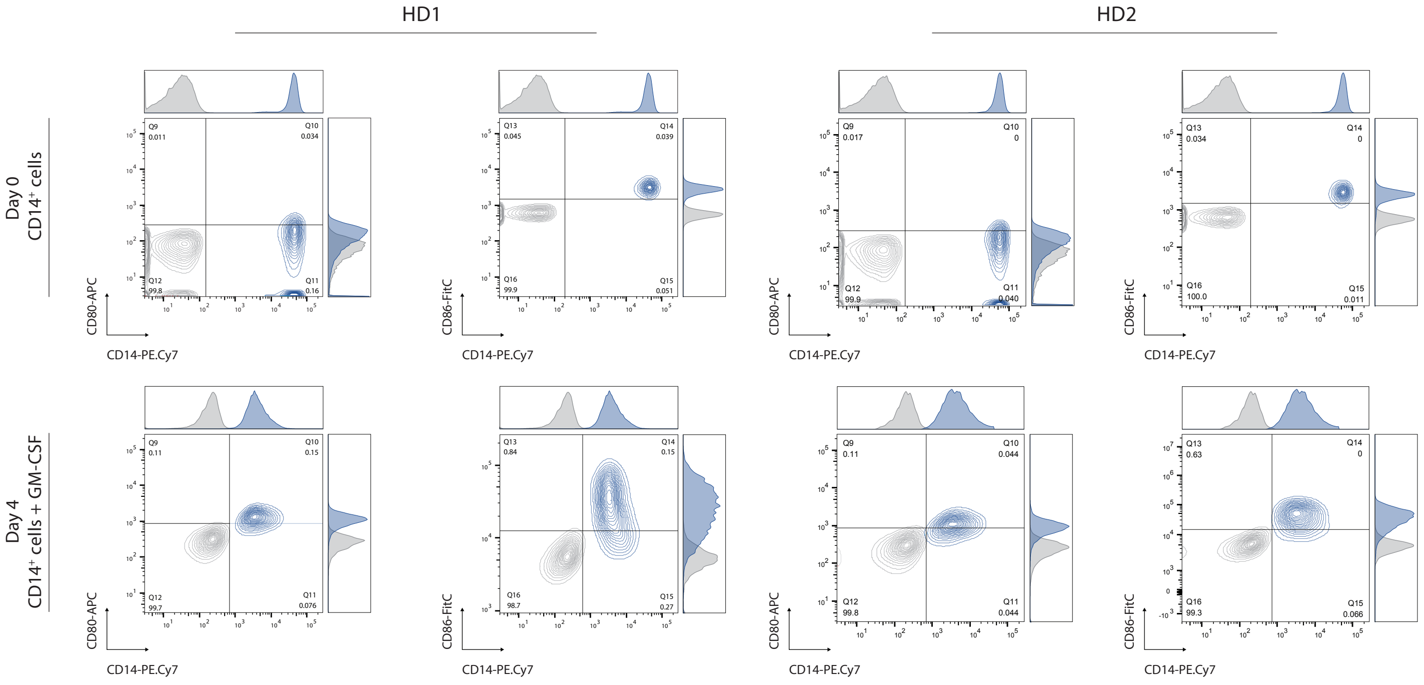

### Supplementary Figure 5

#### Supplementary Figure 5

A

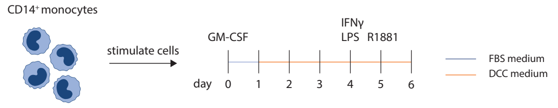


B

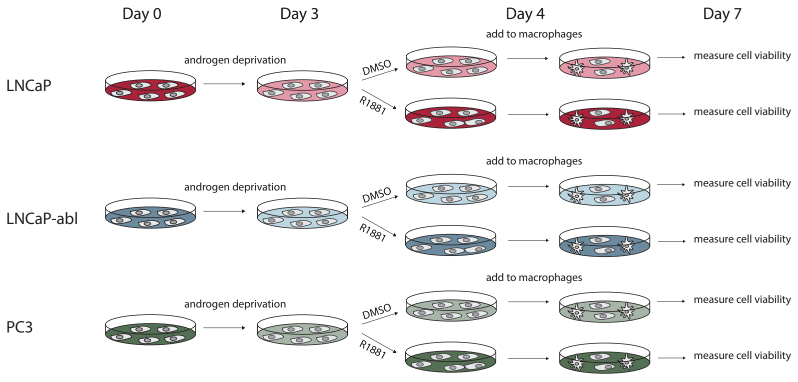


C

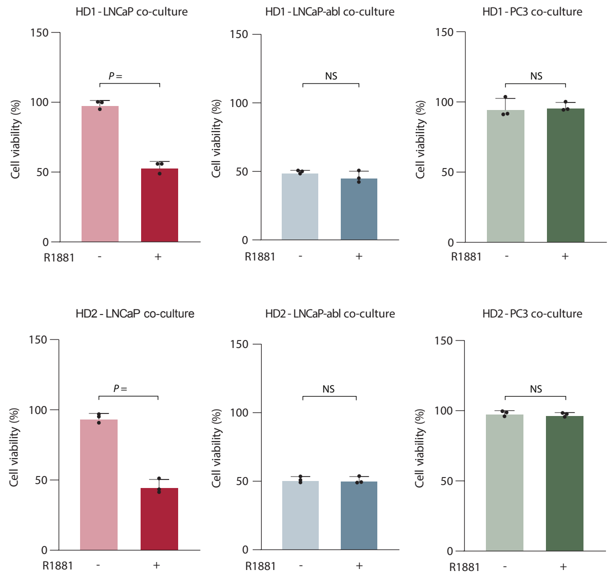

### Supplementary Figure 6

Supplementary Figure 6

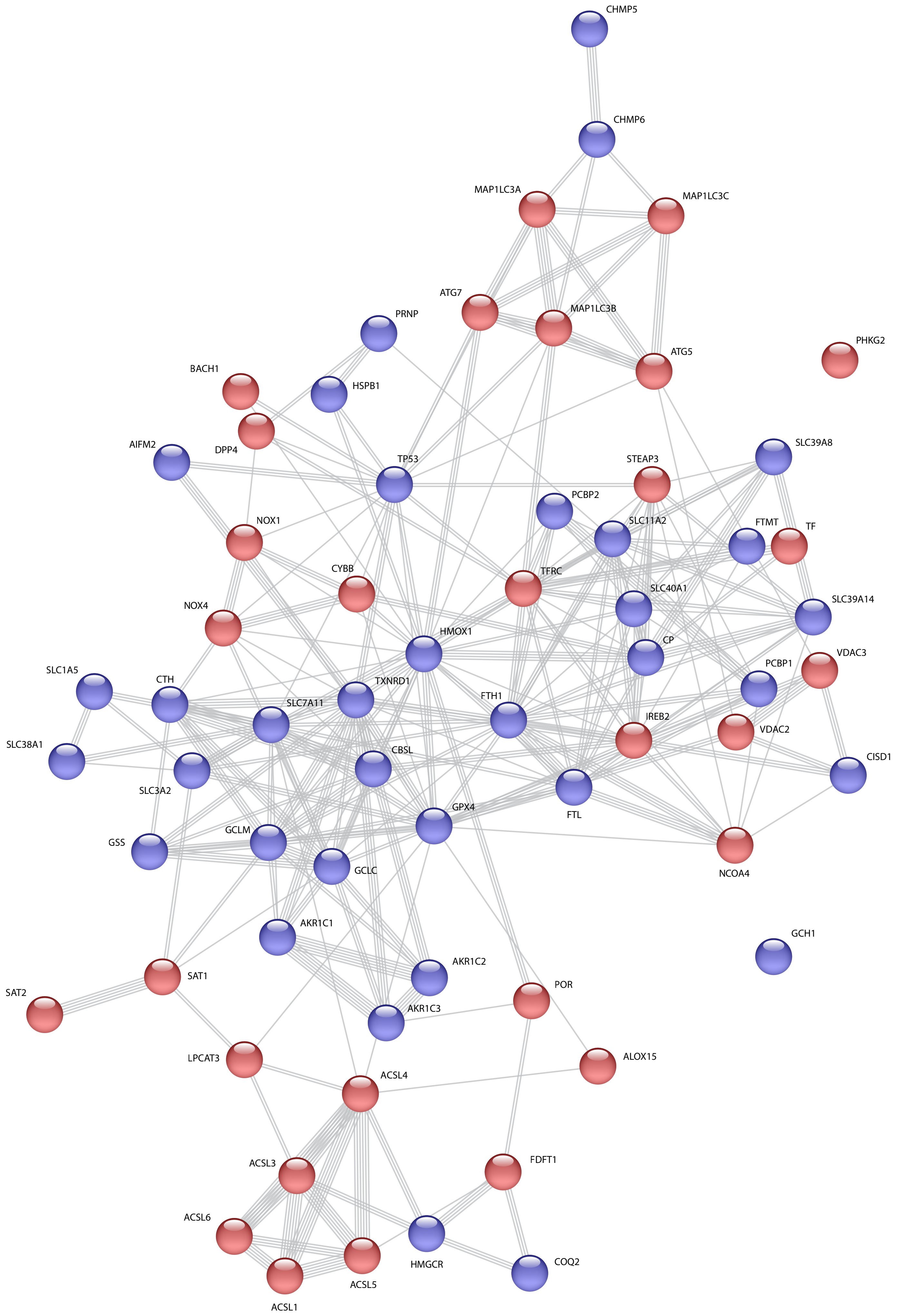

### Supplementary Figure 7

#### Supplementary Figure 7

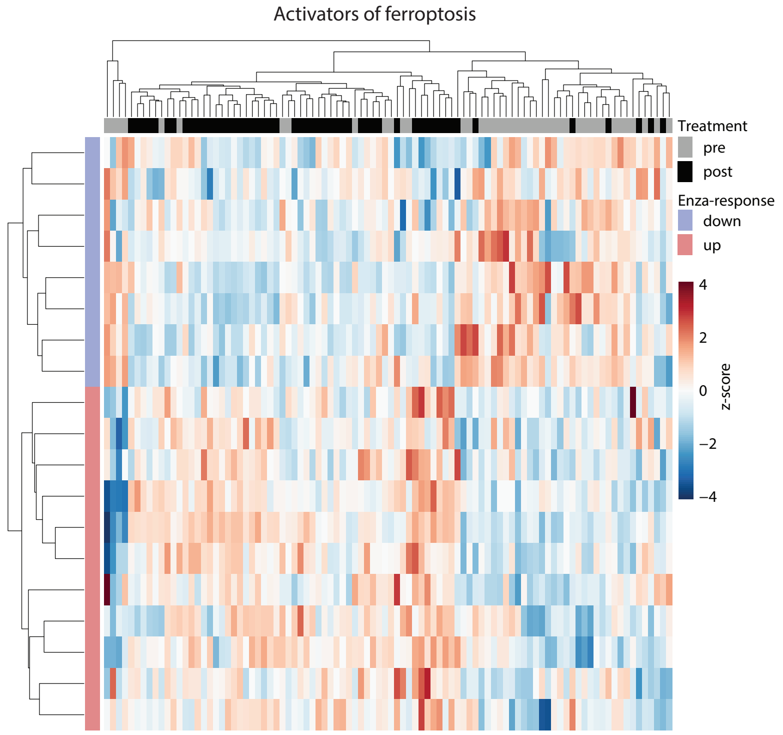
